# Legume plant defenses and nutrients mediate indirect interactions between soil rhizobia and chewing herbivores

**DOI:** 10.1101/2021.11.10.468162

**Authors:** Saumik Basu, Benjamin W Lee, Robert E Clark, Sayanta Bera, Clare L Casteel, David W. Crowder

**Affiliations:** Department of Entomology, Washington State University, Pullman, WA, USA, 99164; School of Integrative Plant Science, Plant Pathology and Plant-Microbe Biology Section, Cornell University, Ithaca, NY, USA, 14850; Department of Cell Biology and Molecular Genetics, University of Maryland College Park, College Park, MD 20742, USA

**Keywords:** pea leaf weevil, defense genes, phytohormones, physical defense, plant nutrients

## Abstract

Soil bacteria that form mutualisms with plants, such as rhizobia, affects susceptibility of plants to herbivores and pathogens. Soil rhizobia also promote nitrogen fixation, which mediates host nutrient levels and defenses. However, whether aboveground herbivores affect the function of soil rhizobia remains poorly understood. We assessed reciprocal interactions between *Sitona lineatus*, a chewing herbivore, and pea (*Pisum sativum*) plants grown with or without rhizobia (*Rhizobium leguminosarum* biovar *viciae)*. We also examined the underlying plant-defense and nutritional mechanisms of these interactions. In our experiments, soil rhizobia influenced feeding and herbivory by chewing herbivores. Leaf defoliation by *S. lineatus* was lower on plants treated with rhizobia, but these insects had similar amino acid levels compared to those on un-inoculated plants. Plants grown with soil rhizobia had increased expression of gene transcripts associated with phytohormone-mediated defense, which may explain decreased susceptibility to *S. lineatus*. Rhizobia also induced expression of gene transcripts associated with physical and antioxidant-related defense pathways in *P. sativum*. Conversely, *S. lineatus* feeding reduced the number of root nodules and nodule biomass, suggesting a disruption of the symbiosis between plants and rhizobia. Our study shows that aboveground herbivores can engage in mutually antagonistic interactions with soil microbes mediated through a multitude of plant-mediated pathways.

## Introduction

Soil harbors abundant and diverse microbe communities that affect ecosystem functions like biomass production, carbon sequestration, pollution mitigation, and nutrient cycling (A’Bear, Johnson & Jones, 2014; Bardgett & van der Putten, 2014). Plant-root associated soil bacteria such as rhizobia can also affect plant susceptibility to herbivores and pathogens by altering plant nutrient levels or physical and chemical defenses (Dean, Mescher & De Moraes, 2014; Rashid & Chung, 2017; Heinen, Biere, Harvey & Bezemer, 2018; Blundell et al., 2020). By affecting plant traits, soil microbes often may indirectly alter interactions between plants, herbivores, and plant pathogens, and integrating aboveground and belowground interactions is a key priority in food web ecology (Hooper et al., 2005; Pangesti, Pineda, Pieterse, Dicke & Van Loon, 2013; van Geem et al., 2013; de Vries & Wallenstein, 2017; Ramirez et al., 2018).

Soil microbes have cascading bottom-up impacts on aboveground organisms by altering plant traits (Pineda, Soler, Pozo, Rasmann & Turlings, 2015; Tao, Hunter & de Roode, 2017; Valencia et al., 2018). For example, legume plants grown in soil inoculated with rhizobia have greater biomass than plants grown without rhizobia as well as greater systemic resistance against herbivores and pathogens (Gopalakrishnan et al., 2015). Soil rhizobia may also indirectly affect herbivores and pathogens by altering plant defense signaling, release of volatile organic compounds, and plant nutrients (Rasmann, Bennett, Biere, Karley & Guerrieri, 2017; Tao et al., 2017; Heinen et al., 2018). Similarly, aboveground pathogens and herbivores may often disrupt plant-microbe mutualisms, resulting in reduced biological nitrogen fixation and weakened plant defense (Heath & Lau, 2011; Ballhorn, Younginger & Kautz, 2014; Simonsen & Stinchcombe, 2014). These studies suggest reciprocal interactions between microbes, herbivores, and pathogens may often be mediated *via* plant-mediated interactions. Yet, few studies have extensively characterized the mechanistic chemical, physical, and nutritional properties of plants and how they may mediate interactions between aboveground and belowground organisms.

Direct herbivore-soil microbe interactions may also occur when herbivores spend part of their life belowground. For example, *Sitona lineatus* (pea leaf weevil) larvae consume nodules of legume roots that harbor rhizobia. However, the majority of interactions between soil microbes and aboveground organisms are likely to be indirect and plant-mediated. For example, rhizobia-inoculated legumes are often less susceptible to herbivory, as physical defenses such as greater callose deposition and induction of antioxidants are promoted in these rhizobia-inoculated plants (Millet et al., 2010; Cawoy et al., 2014; Rashid, Khan, Hossain & Chung, 2017). On the other hand, herbivory may interfere with legume-rhizobia symbiosis, reducing the number and size of root nodules, if aboveground herbivores decrease photosynthesis and plant vigor (Simonsen & Stinchcombe, 2014; Heath & Lau, 2011). By limiting nodule growth and rhizobia function, herbivores might benefit by interfering with anti-herbivore defense signaling induced by rhizobia (Pineda, Zheng, van Loon, Pieterse & Dicke, 2010; Shikano, Rosa, Tan & Felton, 2017; Heinen et al., 2018).

Here we addressed the mechanisms driving trait-mediated indirect interactions between a legume host (*Pisum sativum*, pea), soil rhizobia (*Rhizobium leguminosarum* biovar. *viciae)*, and a chewing herbivore (*S. lineatus)*. In the Palouse region of northern Idaho and eastern Washington, USA, these organisms commonly co-occur in natural and managed ecosystems. However, it is largely unknown if *S. lineatus* herbivores are affected by the presence of rhizobia in soil, or whether herbivory from *S. lineatus* affects symbioses between rhizobia and pea plants. We used greenhouse experiments to assess whether *S. lineatus* affected soil rhizobia and how soil rhizobia affected plant susceptibility to *S. lineatus*. These experiments were complemented with molecular assays that examined chemical, physical, and antioxidant defense signaling and nutritional properties of *P. sativum* hosts exposed to *S. lineatus* and inoculated with *Rhizobium*.

## Materials and Methods

### Study system and experimental conditions

Many native and cultivated legumes, including *P. sativum*, are found in Palouse region of eastern Washington and northern Idaho, USA (Clement, Husebye & Eigenbrode, 2010; Chisholm, Eigenbrode, Clark, Basu & Crowder, 2019). These plants are attacked by insect vectors, pathogens, and chewing herbivores such as *S. lineatus* (Chisholm et al., 2019; Basu, Clark, Bera, Casteel & Crowder, 2021b). *Sitona lineatu*s adults overwinter outside of *P. sativum* fields and migrate into fields in the late spring to lay eggs (Carcamo et al., 2018). After eggs hatch, larvae burrow into the soil to feed and pupate before adults re-emerge in the summer (Carcamo et al., 2018). Thus, *S. lineatus* populations attack *P. sativum* hosts above- and belowground for several months. While *S. lineatus* larvae feed on legume roots belowground, directly affecting the abundance of rhizobia, we focused on adults feeding aboveground to isolate plant-mediated mechanisms by which rhizobia affected this herbivore (Mutch & Yang, 2004).

Adult *S. lineatus* were collected from *P. sativum* fields one wk before experiments, and soil was collected from the Palouse Conservation Farm (Pullman, WA, USA) before being exposed to treatments. For rhizobia treatments, soil was inoculated with pea-specific rhizobia (*Rhizobium leguminosarum* biovar. *viciae)* by mixing N-Dure^R^, a peat-based inoculant with *P. sativum* seeds using the manufacturer’s protocol (Verdasian Life Sciences, Cary, NC, USA). All experiments were conducted in greenhouses at Washington State University (Pullman, WA, USA) with a 16:8 h light:dark cycle, 21-24°C during light cycles, and 16-18°C during dark cycles.

### Effects of rhizobia on S. lineatus feeding

We assessed effects of rhizobia on *S. lineatus* feeding with three soil treatments: (i) control (no treatment); (ii) autoclaved to remove microbes; and (iii) autoclaved with rhizobia added. In autoclaved treatments, field-collected soil was placed in 61 × 91 cm bags in a steam autoclave at 7 psi and 111°C overnight. As autoclaving soil affects soil moisture, all soil treatments were standardized to 75% moisture before plants were added.

Plants were grown in potting mix (Sunshine® LC1) before transplantation into treated soil at 2 wk old. Plants were then individually placed in 1 L pots with soil exposed to one treatment, which were placed in bucket cages (0.6 × 0.3 × 0.3m) for an additional 2 wk before *S. lineatus* treatments were applied. There were two *S. lineatus* treatments: (i) none (control) and (ii) two adult *S. lineatus* feeding on plants for 48 h. After 48 h, both adult *S. lineatus* were removed from each plant to prevent further feeding. The experiment was a 3 × 2 factorial design, with 3 soil treatments and 2 *S. lineatus* treatments; each treatment was replicated 10 times per block, and two temporal blocks were performed. There were a total of 120 experimental units (2 blocks × 3 soil treatments × 2 *S. lineatus* treatments × 10 replicates). In each replicate, the total numbers of leaf notches were counted by visually observing the aboveground portion of all the plants. Leaf notches is a reliable indicator of the amount of *S. lineatus* feeding (Chisholm, Sertsuvalkul, Casteel & Crowder, 2018).

### Analyses of amino acids

We measured amino acid content of *S. lineatus* adults from the different treatments to assess herbivore nutrient acquisition. Two adult *S. lineatus* were collected from each replicate of the feeding experiment (4 replicates per each soil treatment) into liquid N_2_ and lyophilized. After lyophilization, *S. lineatus* tissue was weighed and extracted with 20mM of HCL (Patton, Bak, Sayre, Heck & Casteel, 2019). Amino acids were derivatized using AccQ-Fluor reagent kits (Waters, Milford, MA, USA), with L-Norleucine as an internal standard. 10 μl from each sample were injected into a Agilent 1260 Infinity HPLC (Agilent, Santa Clara, CA, USA) with a Nova-Pak C18 column (c).

Amino acid derivatives were detected with excitation and emission wavelengths of 250 nm and 395 nm, respectively. Peak areas were compared to a standard curve made from a serial dilution of amino acid standards (Sigma-Aldrich, St. Louis, MO). Solvent A, AccQ-Tag Eluent A, was premixed from water; Solvent B was acetonitrile:water (60:40). The gradient used was 0–0.01 min, 100% A; 0.01–0.5 min, linear gradient to 3% B; 0.5–12 min, linear gradient to 5% B; 12–15 min, linear gradient to 8% B; 15–45 min, 35% B; 45–49 min, linear gradient to 35% B; 50– 60 min, 100% B. The flow rate was 1.0 ml min^−1^. Amino acid derivatives and peak areas were measured with an Agilent fluorescence detector and ChemStation software. To calculate concentrations, standard curves were created for each amino acid using dilutions of standards.

### Effects of S. lineatus on soil rhizobia

We next assessed how *S. lineatus* feeding affected nodulation and nodule biomass, two key metrics of rhizobia function, with two treatments: (i) control - no *S. lineatus* and (ii) *S. lineatus* feeding. In *S. lineatus* treatments, we released two adults for 48 h on 2 wk old *P. sativum* plants, after which the adults were removed. Following treatments, plants were uprooted from the soil after 7 d and soil was washed off roots with tap water. Nodules were counted from the root of each plant and then excised. Nodule fresh weights were taken immediately after collection, then dried for 5 d at 37°C before dry weight measurements were taken. Plants that failed to develop any root nodules served as a control for this experiment (no rhizobia inoculation).

### Analyses of transcripts related to defense signaling

We next conducted an experiment with two *S. lineatus* treatments: (i) control, no *S. lineatus* and (ii) two adult *S. lineatus* feeding for 48 h. These treatments were crossed with two rhizobia treatments: (i) control, no rhizobia inoculum and (ii) seeds treated with rhizobia. For preparation of potting mix, soil and sand were mixed in equal volume (1:1) to facilitate nodule development; plants were in treated soil for 2 wk before *S. lineatus* treatments. Plant tissue samples were harvested 3 d and 7 d after *S. lineatus* addition. In total, the experiment included four randomly assigned replicates of each treatment for two temporal blocks in a 2 × 2 × 2 factorial design (2 soil treatments × 2 *S. lineatus* treatments × 2 time points × 4 replicates = 32 total experimental units). Aboveground harvested plant tissue was wrapped in aluminum foil, frozen in liquid N_2_, and kept on dry ice before storing in −80 °C. Samples were ground using a mortar and pestle in liquid N_2_, and 50 to 100 mg of tissue was used for total RNA extraction using Promega SV total RNA isolation kits (Promega, Madison, WI) and cDNA from 1 μg of total RNA using Bio-Rad iScript cDNA synthesis kits. Gene-specific primers (Table S1) were used in qRT-PCR reactions (10 μl) containing 3 μl of ddH2O, 5 μl of iTaq Univer SYBR Green Supermix, 1 μl of primer mix (forward and reverse), and 1 μl of diluted (1:25) cDNA template. The qRT-PCR program had an initial denaturation for 3 min at 95 °C followed by 40 cycles of denaturation at 95 °C for 15 s, annealing for 30 s at 60 °C, and extension for 30 s at 72 °C. For melting curve analysis, a dissociation step cycle was used (55 °C for 10 s, and then 0.5 °C for 10 s until 95 °C). The relative expression of genes were calculated using the delta-delta Ct method, (2 ^−ΔΔCt^) with Psβ-tubulin as a housekeeping gene (Livak & Schmittgen, 2001; Kozera & Rapacz, 2013).

Harvested plant tissue was assessed for expression of 14 gene transcripts associated with hormone signaling, physical, or antioxidant-related defense pathways (Fondevilla, Küster, Krajinski, Cubero & Rubiales, 2011; Tran, You & Barbetti, 2018; Kimura & Kawano, 2015). Gene sequences were obtained using accession numbers of available pea genes or by using the pea marker database (Kulaeva et al., 2017) and blast searching the reference pea genome (Kreplak et al., 2019). We assessed expression of 7 gene transcripts related to phytohormones. *Pathogenesis-related protein 1* (*PR1*) and *Isochorismate synthase1* (*ICS1)* are associated with the salicylic acid (SA) pathway, with *ICS1* involved upstream of SA biosynthesis and *PR1* triggering downstream systemic acquired defenses (Zhang et al., 2010; Fondevilla et al., 2011; Seguel et al., 2018). Two genes, *Lipoxygenase 2* (*LOX2*) and *12-oxophytodienoate reductases 3* (*OPR3*) are associated upstream and downstream, respectively, of jasmonic acid biosynthesis (He, Fukushige, Hildebrand & Gan, 2002; Fondevilla et al., 2011; Wasternack & Hause, 2013). Other genes included were *1-aminocyclopropane-1-carboxylic acid synthases 2* (*ACS2*), which is associated with ethylene biosynthesis, and *Aldehyde oxidase* 3 (*AO3*), which catalyzes abscisic acid biosynthesis. Beside abscisic acid biosynthesis, *AO3* also affects production of reactive oxygen species (Yergaliyev et al., 2016).

We assessed relative transcript accumulation of two additional genes related to physical defense: (i) viz. *β-1,3 Glucanase*, an enzyme that regulates callose production and (ii) *calcium-regulated/ATP-independent ferisome protein gene*, which is associated with P protein plugs that seal phloem pathways (Zavaliev, Ueki, Epel & Citovsky, 2011; Srivastava, Tuteja & Tuteja, 2015; Moravčíková et al., 2016). Six additional genes for antioxidant related defense pathways were assessed: 3 S*uper Oxide Dismutases* (*FeSOD, CuZnSOD, MnSOD*), *Catalase* and *Glutathione reductase 1(GR1*), and *Peroxidase* (*PsPOX11*) (Fondevilla et al., 2011; Tran et al., 2018). Induction of these defense pathways can catalyze superoxides (reactive oxygen species, ROS) in plants (Kimura & Kawano, 2015) and affect induction of salicylic acid in peas (Kawahara et al., 2006).

### Data analysis

Analyses were conducted in R 4.0.5 (R Core Team 2021). We used a generalized linear model (GLM) with a Poisson distribution to assess whether soil treatments (control, autoclaved, rhizobia) affected the number of *S. lineatus* feeding notches; negative controls without *S. lineatus* never had feeding damage, and these treatments were not included. We also used GLMs with a Poisson distribution to assess if *S. lineatus* (present or absent) affected plant nodule weight and nodule biomass. We analyzed effects of soil rhizobia and *S. lineatus* treatments, and their interaction, on fold change gene expression using MANOVA (multiple analysis of variance) on delta CT values (2^−ΔΔCT^ values before transformation) for relative transcript abundance for 14 different genes, *PR1*, *ICS1, OPR3*, *LOX2, AO3*, *ACS2*, *β-1,3 Glucanase, Calcium-regulated/ATP-independent ferisome protein gene, CuZnSOD, FeSOD, MnSOD, Catalase, GR1 and PsOX11*. MANOVA was used as we assumed responses of multiple gene transcripts came from the same plants and therefore have a partially correlated response to treatment. Parameter estimates and subsequent calculations for delta-delta CT (2^−ΔΔCt^) were plotted on the log 10 scale. Finally, average amino acid content for 13 amino acids was fitted to a linear mixed model (LMM, lme4 package, Bates, Maechler, Bolker & Walker, 2015), with soil treatment as a fixed effect and amino acid and replicate as random effects; amino acid concentration was log-transformed to meet normality assumptions. Estimated marginal means and all post-hoc tests were assessed using the emmeans package (Lenth, 2016), with significance tests via analysis of deviance tables generated using the car package (Fox & Weisberg, 2011).

## Results

### Effects of soil rhizobia on *S. lineatus* feeding and amino acid uptake

Rhizobia inoculation altered the amount of herbivory from *S. lineatus* on *P. sativum* hosts (*χ*^2^ = 106.0, *P* < 0.001, Fig. 1). Plants grown in autoclaved soil had the most feeding, while plants grown in autoclaved soil with rhizobia had the least, with plants grown in control soil having intermediate levels (Figs. 1, S1). Soil treatment, however, did not significantly affect the concentration of amino acids in *S. lineatus* from plants (χ^2^ = 1.66, df = 2, *P* = 0.44, Fig. 1B).

**Figure 1.**
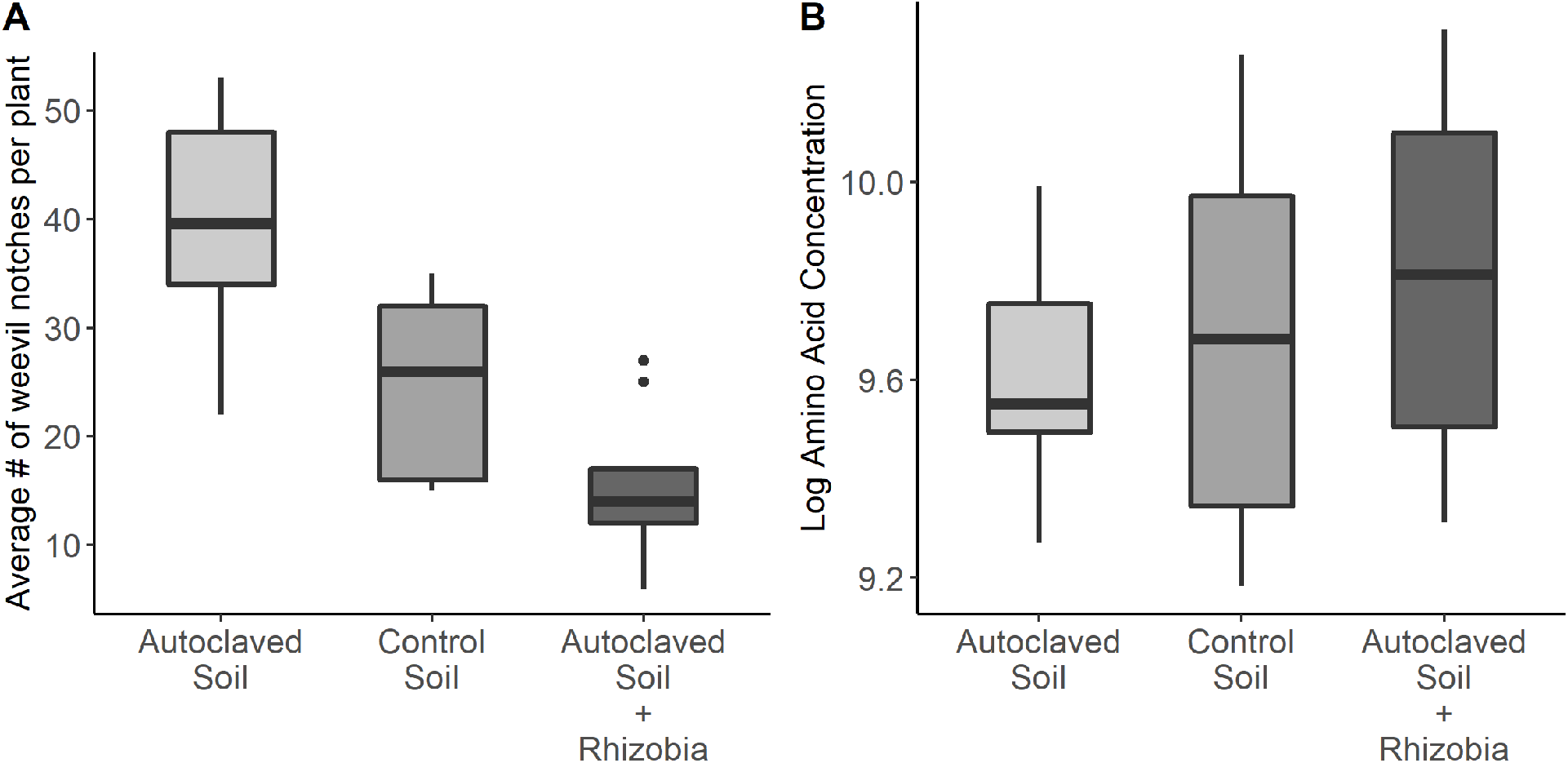
Soil rhizobia confer resistance against *S. lineatuus* feeding. (A) Reduced number of *S. lineatus* induced feeding notches were observed in pea leaves in presence of soil rhizobia (N = 10). Average number of feeding notches in response to soil treatments (Poisson GLM). Rhizobia addition reduced feeding notches, with intermediate levels of herbivory in control soils (p<.05, Tukey HSD). (B) Effect of soil treatments on uptake of amino acids by *S. lineatus*. Log-transformed mean concentrations (nmol/mg DW) among 13 amino acids in weevils feeding on pea plants undergoing various soil treatments.

### Effects of *S. lineatus* herbivory on soil rhizobia

Herbivory from *S. lineatus* had a negative effect on symbiosis between rhizobia and plant hosts (Figs. 2, S2). Plants that were fed on by *S. lineatus* had fewer root nodules (*χ*^2^ = 6.49, *P* = 0.010, Fig. 2A) and lower total nodule biomass (*χ*^2^ = 9.41, *P* = 0.002, Fig. 2B) than plants that did not experience herbivory. However, treatments with *S. lineatus* did not significantly affect nodule dry mass (*χ*^2^ = 2.46, *P* = 0.12, Fig. 2C).

**Figure 2.**
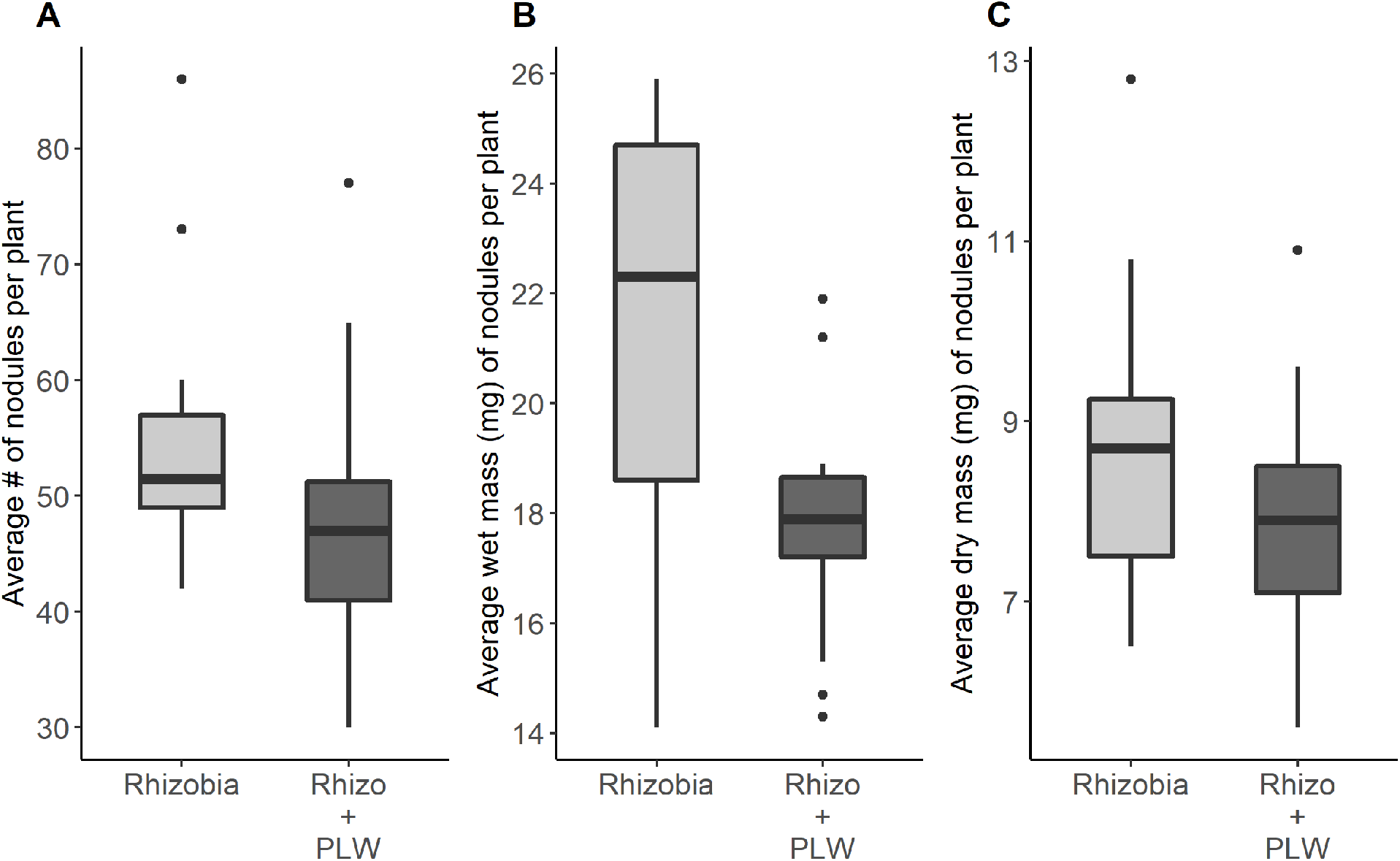
Effect of *S. lineatus* herbivory on nodulation: Fig 2A, B & C. Nodule count based on Poisson-fit GLM. Nodule wet and dry mass based on Gaussian-fit GLM. *S. lineatus* feeding negative affects nodule number and wet weight but not dry mass (p<0.05, Tukey HSD).

### Effects of *S. lineatus* and soil rhizobia on expression of defense gene transcripts

Plants grown with rhizobia had higher expression of the ethylene biosynthetic gene, *ACS2*, compared to plants grown in control soil with or without *S. lineatus* (Fig. 3A). The presence of *S. lineatus* affected expression of other gene transcripts when plants were grown with rhizobia. Plants grown with rhizobia and no herbivory had greater expression of two gene transcripts associated with jasmonic acid, *OPR3* and *LOX2*, and one associated with the final step of abscisic acid biosynthesis, *AO3*, compared to plants grown with rhizobia but no herbivory (Figs. 3B, D, E). *Sitona lineatus* increased levels of the salicylic acid-associated gene transcript, *ICS1*, on control plants compared to plants grown with rhizobia (Fig. 3C). Soil rhizobia also strongly induced *β-1,3 glucanase*, which is associated with callose-mediated defense (Fig. 4A, S3).

**Figure 3.**
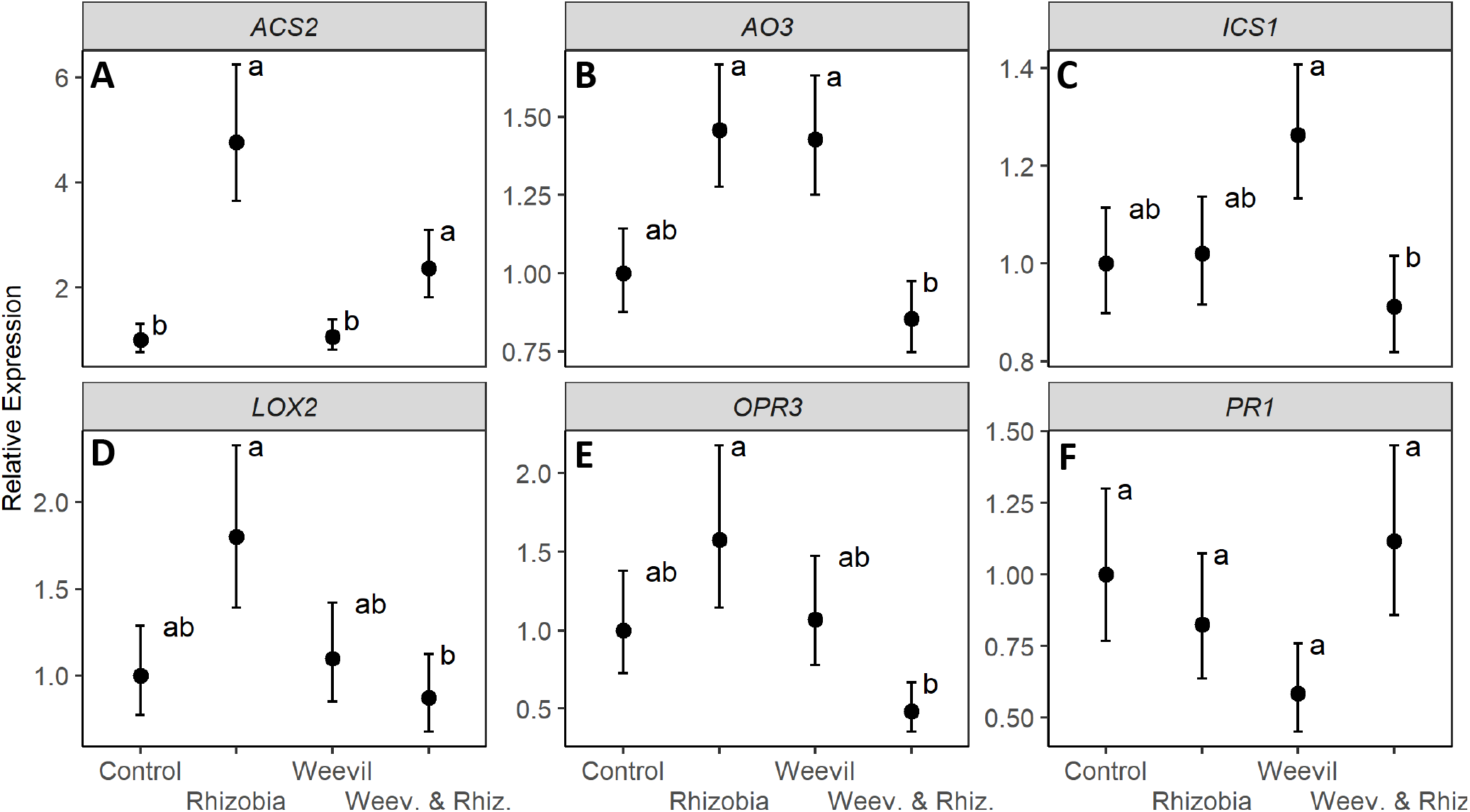
Relative transcript accumulation of SA responsive genes: *ICS1* (A), *PR1* (B); ABA responsive gene: *AO3* (C); JA responsive genes: *OPR3* (D), *LOX2* (E); Ethylene responsive genes: *ACS2* (F) in *Pisum sativum* at 7dpi. Bars not connected with same number are significantly different.

**Figure 4.**
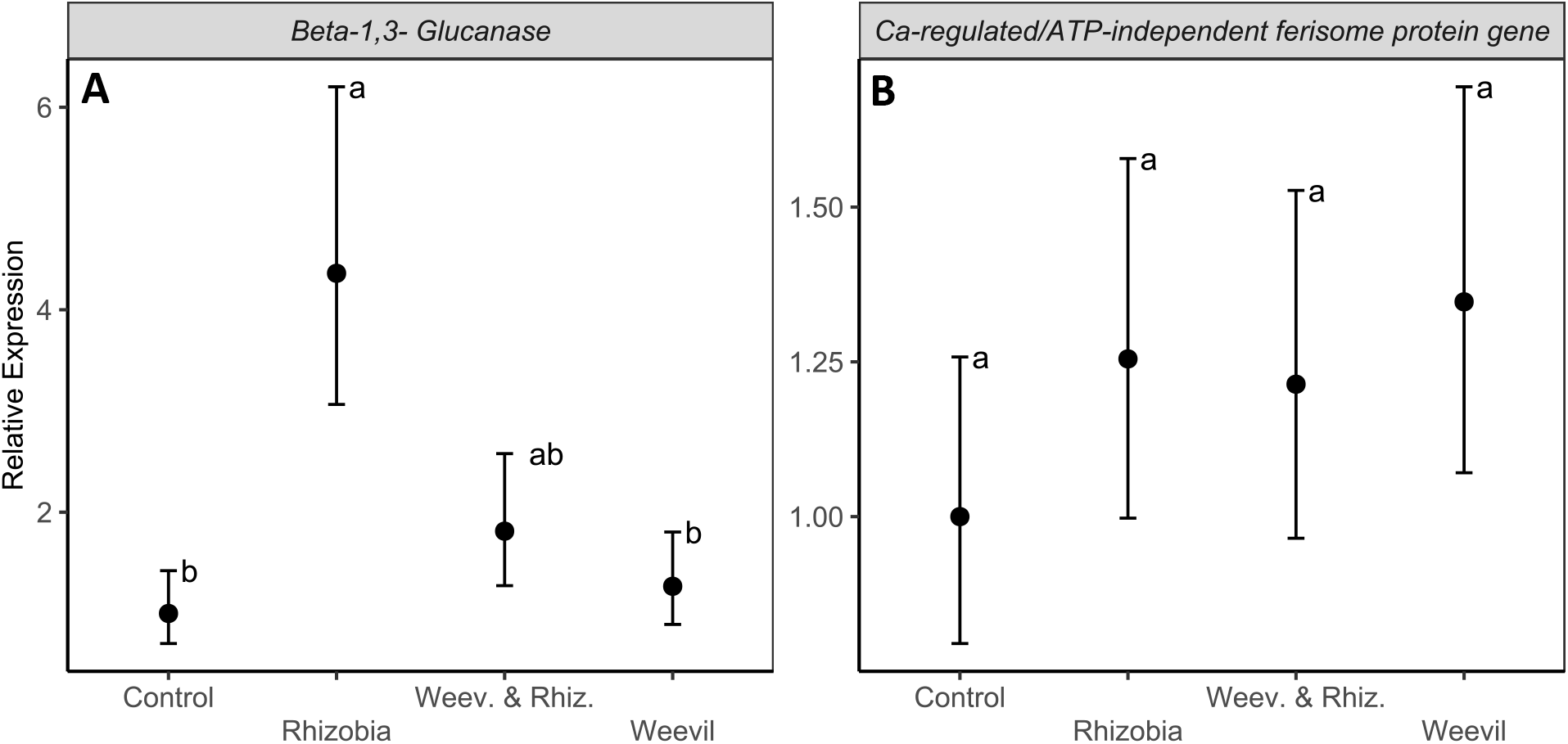
Relative transcript accumulation of callose mediated defense genes: *Beta-1, 3 glucanase* (A), *PR1* (B) and *Calcium-regulated/ATP-independent ferisome protein gene* in *Pisum sativum* 7dpi. Bars not connected with same number are significantly different.

For the six gene transcripts related to antioxidant-mediated defense, three genes associated with the superoxidase disumaste (*FeSOD*, *CuZnSOD*, *MnSOD*) had greater expression in plants grown in rhizobia-inoculated soil compared to control soil (Figs. 5B, C, E, S3). However, other gene transcripts were not impacted by rhizobia. We found that *S. lineatus* induced the antioxidant gene transcript, *Catalase*, on plants grown in soil without rhizobia (Figs 5A, S3), and the gene transcript peroxidase (*PsPOX11*) on plants grown in soil with rhizobia (Figs. 5F, S3).

**Figure 5.**
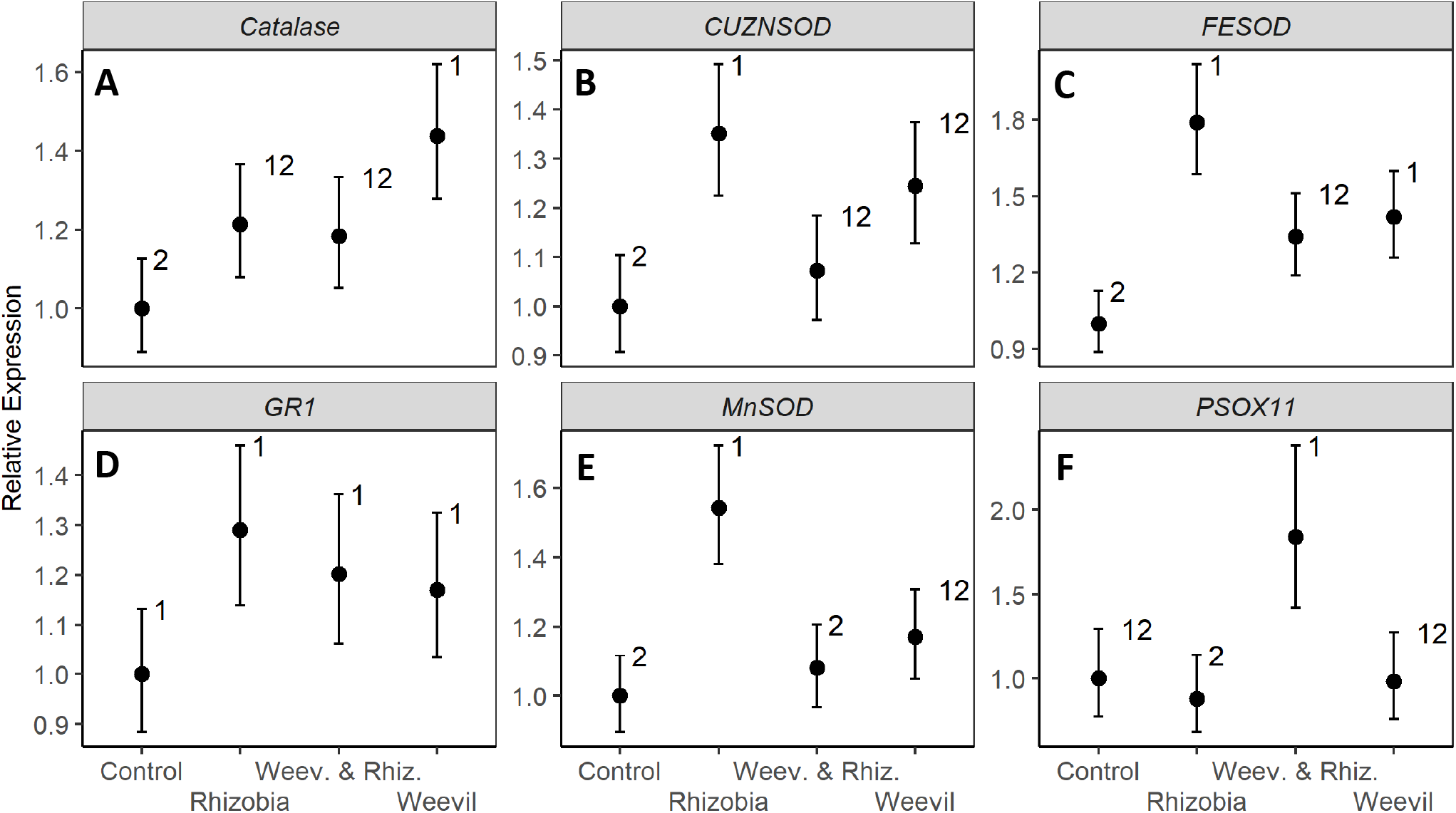
Relative transcript accumulation of antioxidant related defense genes: *CuZnSOD* (A), *FeSOD* (B), *MnSOD*(C), *Catalase* (D), *GR1* (E) and Peroxidase, *PsOX11* (F) in *Pisum sativum* at 7 dpi Bars not connected with same number are significantly different.

## Discussion

Herbivores and soil microbes interact through many direct and indirect, trait-mediated, pathways. Our study highlights plant-mediated mechanisms that may underlie these interactions. Root-associated bacteria can alter transcript levels of important anti-herbivore defensive genes in host plants, which can impact the nutritional quality of plants and nutrient uptake by herbivores. In turn, these changes led to reduced leaf herbivory on plants. We observed that *P. sativum* host plants grown in rhizobia-inoculated soil had reduced leaf defoliation from *S. lineatus* compared to plants grown without rhizobia (either control or autoclaved). However, *S. lineatus* individuals obtained similar levels of amino acids on plants grown in rhizobia-inoculated and control plants. This shows weevils obtained more nutrients per unit of leaf area on plants grown in rhizobia-inoculated soil, which may be due to improved nutritional quality of the host plants (Kempel, Schädler, Chrobock, Fischer & van Kleunen, 2011). Our results raise the intriguing possibility that mutualistic soil microbes promote plant health by increasing plant nutrients, which in turn reduces total feeding by herbivores.

Our results are in line with studies showing rhizobia and arbuscular mycorrhizal fungi are keystone microbes that decrease plant susceptibility to insects and pathogens (Jaber & Vidal, 2009; Pineda et al., 2010; Santos et al., 2014; Yang et al., 2014; Gopalakrishnan et al. 2015; Mabrouk et al., 2018). Our study provides evidence that several mechanisms may underlie these results. Soil rhizobia can modify plant nutrients in legumes as well as various defense related signaling pathways against insects, which can alter insect feeding responses and performance (Dean et al., 2014). For example, soil rhizobia increase tolerance of soybean plants (*Glycine max*) to soybean aphid (*Aphis glycine*), although different rhizobia strains vary in their effects (Dean et al., 2014). Similarly, growth and performance of cotton leaf worm (*Spodoptera littoralis*) is limited by rhizobia on clover (*Trifolium repens*), as rhizobia increased production of nitrogen-based defense compounds in hosts (Kempel, Brandl & Schädler, 2009). However, these effects were not observed on clover plants that were naturally cyanogenic, suggesting benefits of rhizobia may only occur on plants that are not naturally well defended (Kempel et al., 2009).

Rhizobia-induced changes in phytochemical and nutritional traits of plants have increasing been recognized as important drivers of ecosystem function in multi-trophic food webs (Qchieno et al., 2021). For example, beneficial root colonizing soil rhizobia often elicit induced systemic resistance against insects and pathogens though activation of JA and ET signaling (Romera et al., 2019). Symbiotic association of legume roots by soil rhizobia has been associated with enhanced resistance against aboveground consumers including beetles (Soundararajan, Chitra, & Geetha, 2013; Thamer, Schädler, Bonte & Ballhorn, 2011; Godschalx, Tran, & Ballhorn, 2017). Conversely, aboveground feeders damage leaves through defoliation and interfere with photosynthesis by consuming sugars and other nutrients that are required for root nodulation (Katayama et al., 2014). We observed plants attacked by *S. lineatus* had fewer nodules and lower nodule biomass than controls, suggesting antagonistic effects of *S. lineatus* on legume-rhizobia symbiosis. Previous studies have also shown that outbreaks of *S. lineatus* can promote the spread of aphid-borne viruses that also impede the function of rhizobia (Chisholm et al., 2019; Basu et al., 2021a); thus, we have shown that *S. lineatus* may negatively affect plant-rhizobia symbiosis through multiple indirect pathways.

Our analysis of phytohormone transcripts suggests interactions between rhizobia and *S. lineatus* were mediated by phytohormones. Rhizobia induced plant defense against *S. lineatus* by activating jasmonic acid, ethylene, and abscisic acid signaling. Jasmonic acid and ethylene are two key systemic defense pathways induced in plants against chewing herbivores (Pangesti et al., 2015; 2016; Rashid & Chung, 2017; Zhu et al., 2018). Similarly, beneficial microbes often also stimulate biosynthesis of abscisic acid (Sgroy et al., 2009; Jha & Subramanian, 2013), even though abscisic acid signaling can have negative effects on nodulation (Tominaga et al., 2010; Roy Choudhury, Johns & Pandey, 2019). Our study provides further evidence that rhizobia can affect herbivores by altering physical defenses such as callose (Ballhorn et al., 2014; Gaudioso-Pedraza et al., 2018) and antioxidants (Walz, Juenger, Schad & Kehr, 2002, Dumanović, Nepovimova, Natić, Kuča, & Jaćević, 2021). Antioxidant mediated defenses are found in cellular organelles such as chloroplasts, mitochondria, and peroxisomes, and we found evidence that these defenses (*MnSOD*, *FeSOD* and *CuZnSOD*) were impacted positively by rhizobia. These results suggest broad induction of both chemical and physical defense by rhizobia can affect herbivores through trait-mediated indirect pathways.

Overall, our study shows soil rhizobia improve plant health by inducing broad-spectrum systemic resistance against herbivores while also improving plant quality. On the other hand, herbivores can interfere strongly with legume-rhizobia symbiosis by inhibiting root nodule development. Thus, assessing reciprocal interactions between soil rhizobia and herbivores is crucial for understanding broader dynamics of agricultural and natural food webs. As legumes are commonly included in rotations with cereals (corn, rice, wheat, barley) in agroecosystems around the world, understanding how soil microbes (e.g. rhizobia) can be affected by chewing herbivores could lead to more effective management of biological nitrogen fixation and crop sustainability. Manipulation of soil microbes, for example, may also provide a novel tactic to manage devastating herbivores while improving crop yield and nitrogen fixation.

## Author Contributions

S.B^1^. and D.W.C. conceived the ideas and methodology; S.B.^1^, B.W.L., R.E.C., S.B^2^. and C.L.C. collected the data; B.W.L., S.B.^1^, R.E.C., S.B^2^., C.L.C and D.W.C. analyzed and interpreted the data; all authors contributed critically to the drafts and gave final approval for publication.

## Acknowledgements

We thank the many undergraduates who helped in various experiments, data collection, and R. Eposado for assistance with data analysis. This research was supported by USDA-NIFA Grants 2016-67011-24693, 2017-67013-26537, and Hatch project 1014754.

