## supplementary file for "Legume plant defenses and nutrients mediate indirect interactions between soil rhizobia and chewing herbivores"

### Table S1: List of primers used for this study

| Gene | Primer sequences | NCBI Accession No. | Amplicon size (bp) |
|---|---|---|---|
| *PsPR1 F*<br>*PsPR1 R* | TGGGGCAGTGGTGACATAAC<br>TGCGCCAAACAACCTGAGTA | LT635896 | 178 |
| *ICS1 F*<br>*ICS1 R* | CTGGAACAGGAATAGTGGAAGG<br>TCAGAGGCAAGTCCAGTTTG | Psat7g237240.1 | 105 |
| *LOX2 F*<br>*LOX2 R* | GCAACCAAGTGACGAAGTCTA<br>GGAGACCCGATTGTAAGGTATTT | PsCam 059875 | 95 |
| *PsOPR3 F*<br>*PsOPR3 R* | GGGTTGAAATCCATGGGGCT<br>GTAATGCAAACCGACAGCGG | AB095740.1 | 117 |
| *PsACS2 F*<br>*PsACS2 R* | GGCATAGTAATTTGAGGTTGAGCC<br>GCCCCAACATTTAAAGGACCTATTA | AF016459 | 103 |
| *PsAO3 F*<br>*PsAO3 R* | TTATAGGACACAGGCTAGCTCAGCA<br>TGACACAAGCTTATTCAGCATGACA | EF491600.1 | 127 |
| *Psβ-1,3 Glucanase F*<br>*Psβ-1,3 Glucanase R* | GATGGAGGCTCTGCAACTTC<br>GTAGCCCAAGGCCTTCTAGG | LT898513.1 | 98 |
| *Psforisome F*<br>*Psferisome R* | GATTTTGTGGATCCCCATTG<br>CCTGTGGCCTGGTAACTCAT | GQ478228.1 | 121 |
| *PsCu/ZnSODII F*<br>*PsCu/ZnSODII R* | GTTCGTATCACTGGCCTTACTC<br>ATGTGGTCCTGTTGAGATACAC | X56435.1 | 93 |
| *PsMnSOD F*<br>*PsMnSOD R* | AAGCTCCAGAATGCCATCAA<br>GTTCACCACCTCCTTCACTAAC | X60170.1 | 97 |
| *PsFeSOD F*<br>*PsFeSOD R* | CTCTTGCAACTGAGGAGGATAAA<br>GGTAAGGAGTGGATGATGATGG | AY426764.1 | 95 |
| *PsCatalase F*<br>*PsCatalase R* | AAGCAGGCTGGAGAAAGATAC<br>GTGGATCGGTGTCGGATAAA | X60169.1 | 94 |
| *PsGR1 F*<br>*PsGR1 R* | AGGCCGTGGAAAGATTGTAG<br>GTCGACCTCCAACTGAAACTAA | X60373.1 | 92 |
| *PsOX11* F<br>*PsOX11* R | CTTGGAGGACCCACATGGAT<br>TTTGGCTTGCTGTTCTTGCA | AB193816.1 | 61 |
| *β-tubulin* FP<br>*β-tubulin* RP | GTAACCCAAGCTTTGGTGATC<br>ACTGAGAGTCCTGTACTGCT | X54844.1 | 203 |

**Supplementary figure S1:** Effect of soil treatments (control, sterilized and rhizobia) on *S. lineatus* mediated leaf defoliation in pea plants (shown are average plants from each treatments). Number of leaf notches produced by *S. lineatus* induced leaf defoliation were reduced following soil rhizobia treatment.

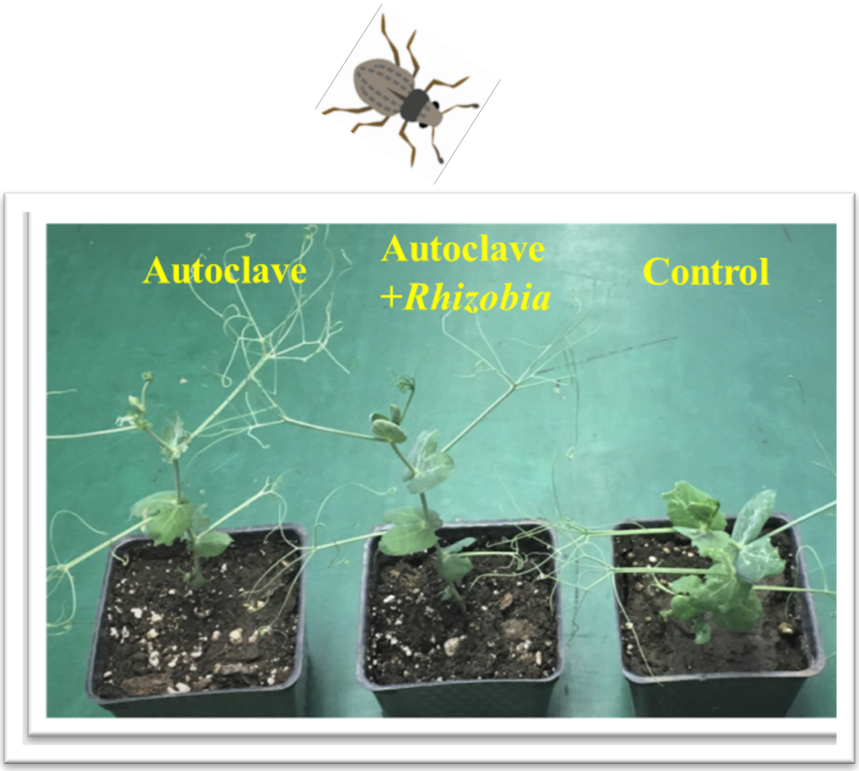

**Supplementary figure S2:** Effects of *S. lineatus* feeding on nodule formation by soil rhizobia (*Rhizobium leguminosarum* biovar. *viciae*) in pea roots. *S. lineatus* feeding reduces nodule no. in (A) intact roots and (B) nodule size as shown in detached nodules.

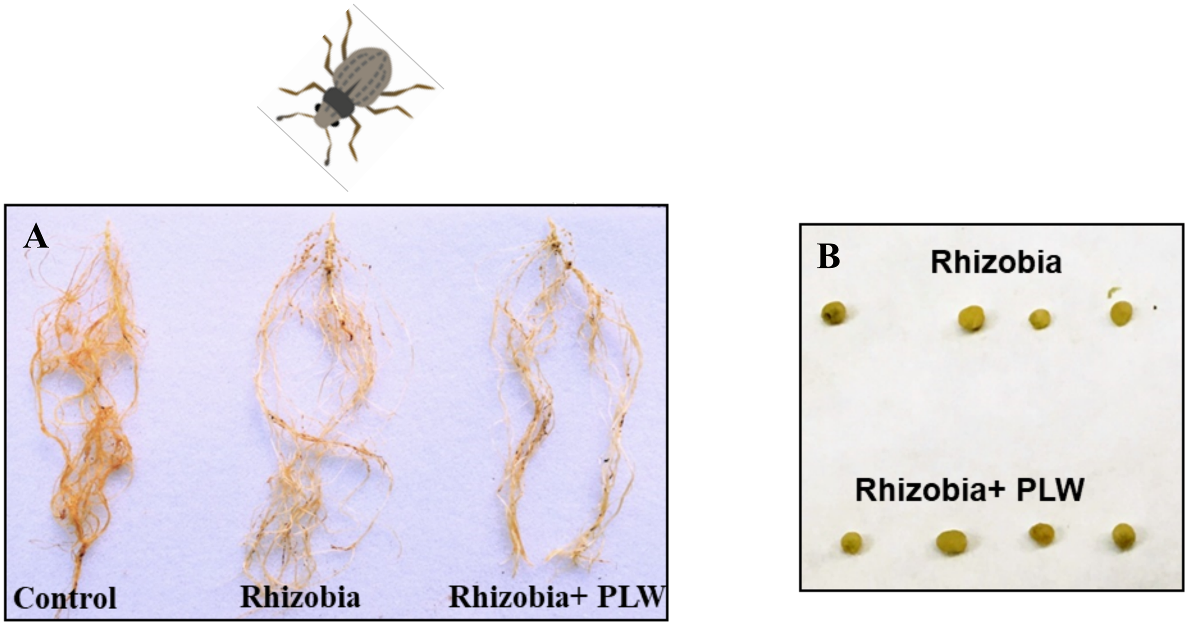

**Supplementary Figure S3:** Relative transcript accumulation of 15 different genes associated with phytohormone signaling, physical defense and antioxidant related defense in *Pisum sativum* at 3 dpi. Bars not connected with same number are significantly different.

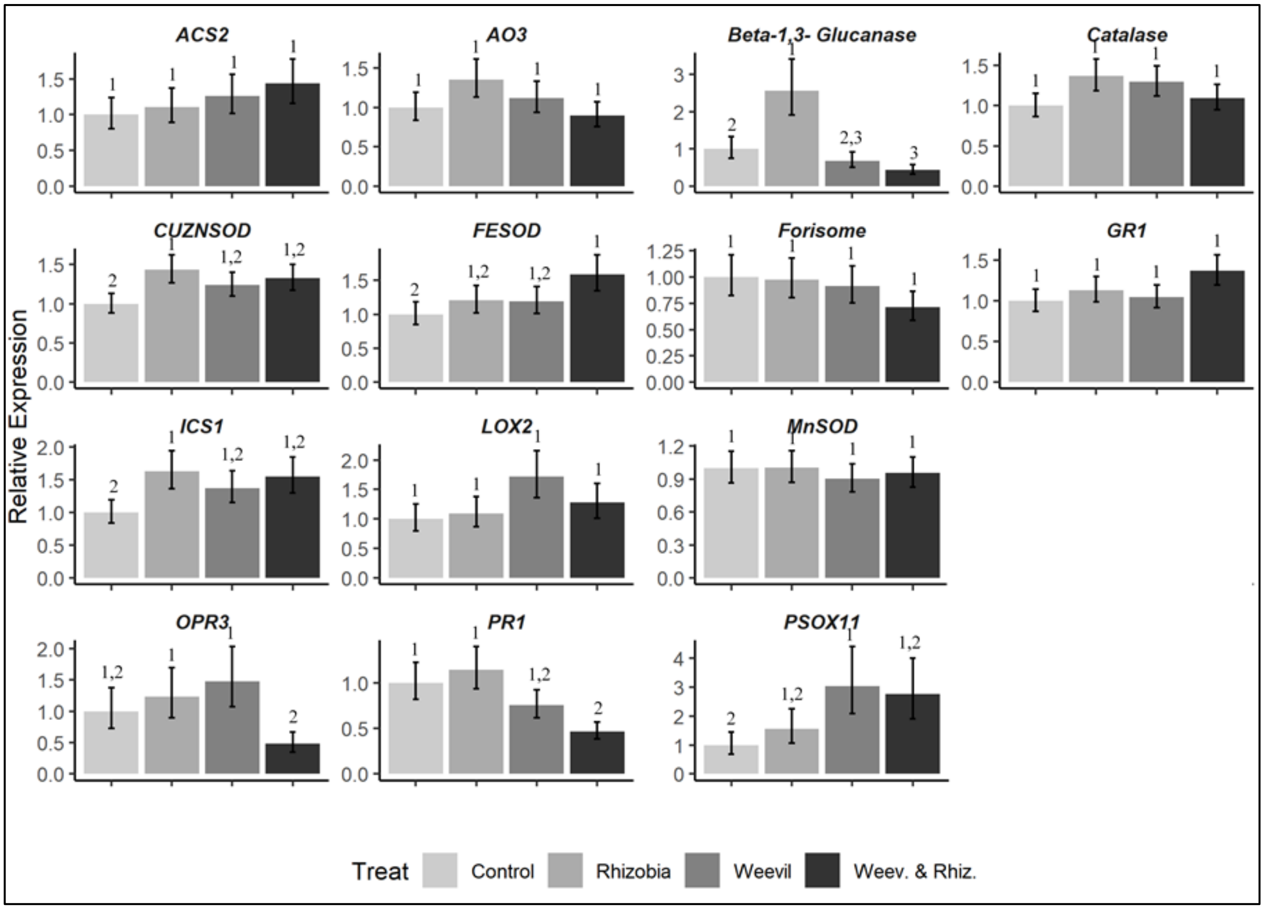

**Supplementary Figure S4:** Effect of soil treatments on uptake of 13 different amino acids by *S. lineatus* from pea plants. Log-transformed amino acid concentrations (nmol/mg DW) separately for 13 amino acids from *S. lineatus* feeding on pea plants undergoing various soil treatments.

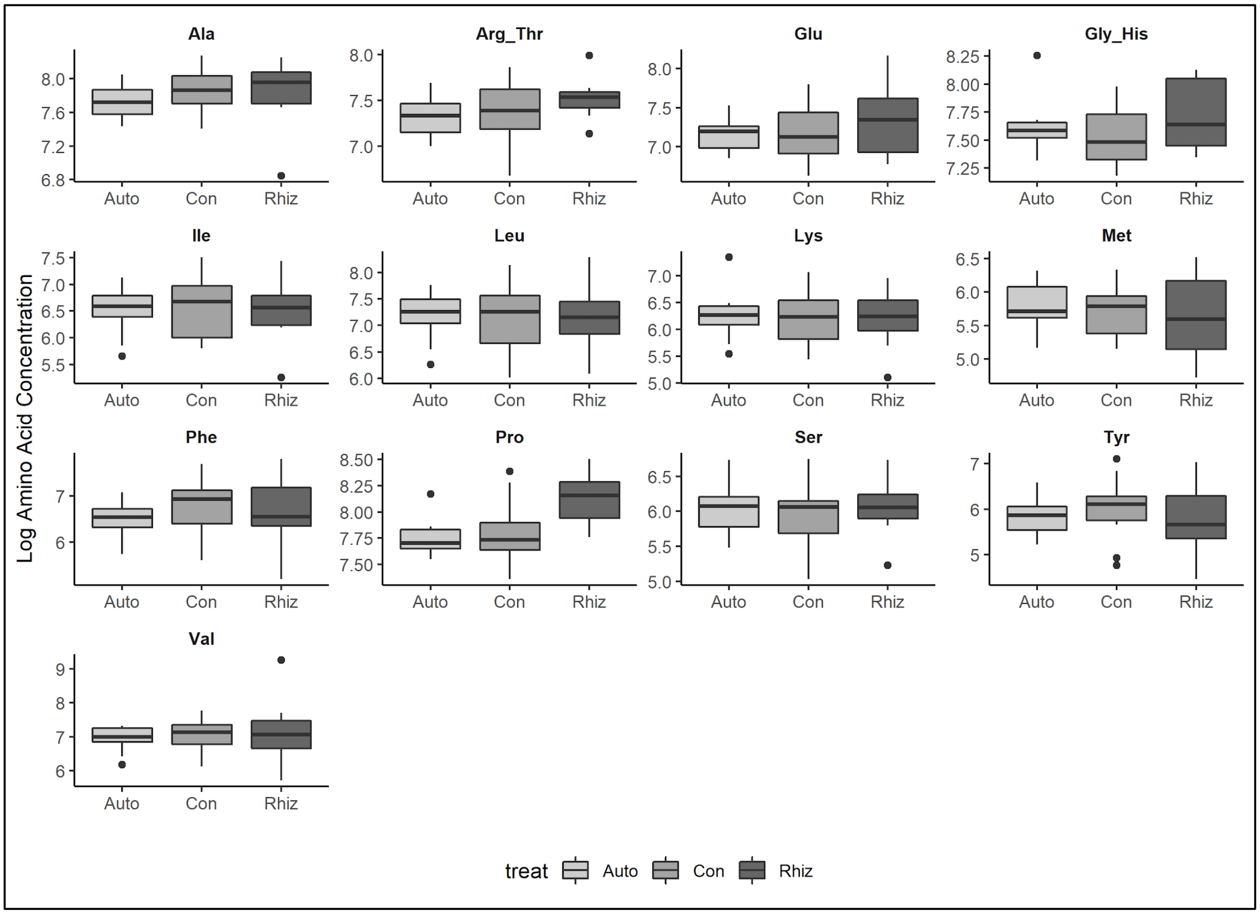
